# The role of parvalbumin interneuron GIRK signaling in regulation of affect and cognition in male and female mice

**DOI:** 10.1101/2020.10.28.359067

**Authors:** EM Anderson, S Demis, H D’Acquisto, A Engelhardt, M Hearing

**Author notes:** **Corresponding Author:** Matthew C Hearing, 560 16^th^ Street, Milwaukee, WI 53233.

## Abstract

Pathological impairments in the regulation of affect (i.e. emotion) and flexible decision-making are commonly observed across numerous neuropsychiatric disorders and are thought to reflect dysfunction of cortical and subcortical circuits that arise in part from imbalances in excitation and inhibition within these structures. Disruptions in GABA transmission, in particular that from parvalbumin-expressing interneurons (PVI), has been highlighted as a likely mechanism by which this imbalance arises, as they regulate excitation and synchronization of principle output neurons. G protein-gated inwardly rectifying potassium ion (GIRK/Kir3) channels are known to modulate excitability and output of pyramidal neurons in areas like the medial prefrontal cortex and hippocampus, however the role GIRK plays in PVI excitability and behavior is unknown. Male and female mice lacking GIRK1 in PVI (Girk1^flox/flox^:PVcre) and expressing td-tomato in PVI (Girk1^flox/flox^:PV^Cre^:PVtdtom) exhibited increased open arm time in the elevated plus maze, while males showed an increase in immobile episodes during the forced swim test. Loss of GIRK1 did not alter motivated behavior for an appetitive reward or impair overall performance in an operant-based attention set-shifting model of cognitive flexibility, however it did alter types of errors committed during the visual cue test. Unexpectedly, baseline sex differences were also identified in these tasks, with females exhibiting overall poorer performance compared to males and distinct types of errors, highlighting potential differences in task-related problem solving. Interestingly, reductions in PVI GIRK signaling did not correspond to changes in membrane excitability but did increase action potential firing at higher current injections in PVI of males, but not females. This is the first investigation on the role that PVI GIRK-signaling has on membrane excitability, action potential firing, and their role on affect and cognition together increasing understanding of PVI cellular mechanisms and function.

## INTRODUCTION

Cognitive flexibility is the ability to adapt behavior in response to changing environmental contingencies and is a critical component to everyday life. As such, impairments in flexibility increase susceptibility to negative life events (e.g., stress), reduce emotional control, and promote development of maladaptive behaviors that can disrupt the capacity of individuals to engage in their lives effectively (Gabrys, Tabri, Anisman, & Matheson, 2018; Lange, Seer, & Kopp, 2017; Waltz, 2017). Neuropsychiatric disorders such as major depression disorder (MDD), obsessive-compulsive disorder (OCD), autism, and schizophrenia share common symptomology including cognitive inflexibility, reduced inhibitory control, and impaired working memory (Dajani & Uddin, 2015; Diamond, 2013; Etkin, Gyurak, & O’Hara, 2013; Marazziti, Consoli, Picchetti, Carlini, & Faravelli, 2010; Moghaddam & Javitt, 2012; Remijnse et al., 2013) however what contributes to these impairments is not well understood.

Optimal regulation of affect (i.e. emotion) and flexible decision-making require a balance of excitation and inhibition (E/I) in numerous cortical and subcortical circuits including the medial prefrontal cortex (mPFC) (Gandal, Nesbitt, McCurdy, & Alter, 2012; Kehrer, Maziashvili, Dugladze, & Gloveli, 2008; Murray et al., 2015; Pantazopoulos, Lange, Hassinger, & Berretta, 2006; Yizhar et al., 2011). Pathological alterations in GABA transmission, in particular that of parvalbumin-expressing interneurons (PVI), has been highlighted as a likely mechanism by which E/I imbalances and associated symptomology arises (Cardin et al., 2009; Ferguson & Gao, 2018; Gandal, Sisti, et al., 2012; Kim, Ahrlund-Richter, Wang, Deisseroth, & Carlen, 2016; Murray et al., 2015; Sohal, Zhang, Yizhar, & Deisseroth, 2009; Wöhr et al., 2015). Analysis of postmortem human brain tissues revealed decreased expression of parvalbumin and parvalbumin mRNA in patients affected by schizophrenia or autism (Curley & Lewis, 2012; Hashemi, Ariza, Rogers, Noctor, & Martínez-Cerdeño, 2017; Hashimoto et al., 2003; Lewis, 2014). Similarly, loss of parvalbumin in rodents promotes autism- (Wöhr et al., 2015) and depression-like symptoms (Fogaça & Duman, 2019), together suggesting a role for parvalbumin, and PVI, in related neuropsychiatric disorder symptomatology.

Within cortical regions such as the mPFC, PVIs are powerful coordinators of network activity (Kepecs & Fishell, 2014; Klausberger & Somogyi, 2008; Markram et al., 2004; Rudy, Fishell, Lee, & Hjerling-Leffler, 2011), with fast-spiking PVIs comprising approximately 50% of cortical interneurons (Kawaguchi & Kubota, 1997). PVIs target the soma and perisomatic compartments of principle output pyramidal neurons where they regulate excitation and synchronize firing (Atallah, Bruns, Carandini, & Scanziani, 2012; Celio & Heizmann, 1981; Ferguson & Gao, 2018; Hu, Gan, & Jonas, 2014; Kubota & Kawaguchi, 1994; Kvitsiani et al., 2013) to orchestrate cortical information flow (H. Kim et al., 2016; Murray et al., 2015; Sohal et al., 2009). In the mPFC, PVIs are highly recruited by afferent excitatory signaling, however intrinsic cellular properties (e.g., membrane excitability) dictate how cells respond to this excitatory drive.

G protein-gated inwardly rectifying potassium ion (GIRK/Kir3) channels produce a slow hyperpolarizing current which modulates neuron excitability and spike firing (Hearing et al., 2013; Marron Fernandez de Velasco et al., 2015; Nimitvilai, Lopez, Mulholland, & Woodward, 2017) acting through inhibitory G protein-coupled receptors including GABA_B_R (Glaaser & Slesinger, 2015; Luján & Aguado, 2015). The role of mPFC and forebrain GIRK channels on pyramidal neuron excitability and behavioral outcomes has been established (Hearing et al., 2013; Victoria et al., 2016), however while GIRKs are known to reside in PVIs and likely contribute to GABA_B_R-mediated signaling in the hippocampus (Booker et al., 2013), their role on mPFC PVI excitability and output is not known. Increasing evidence suggests that drugs that target G protein inhibitory signaling may serve as clinically relevant therapeutic strategies to treat both cognitive and affect-related symptoms (Gandal, Nesbitt, et al., 2012; Kumar et al., 2013; Lecca et al., 2016; Mombereau et al., 2004; Slattery, Desrayaud, & Cryan, 2005). Thus, identifying key modulators of this signaling and how it varies across cell-types remains an important step. Similarly, while PVIs have become a target of recent therapies, there is a need to better understand the cellular mechanisms that regulate their function before progress can be made in this regard (Hu et al., 2014). Accordingly, this study focuses on inhibitory signaling mediated by PVI GIRK channels by determining the role this signaling has on output of PVI in the prelimbic cortex and its relevance to prefrontal cortex-dependent regulation of affect and cognitive flexibility in both male and female mice.

## MATERIALS AND METHODS

### Animals

*Girk1^flox/flox^* stock mice were generated as described (Kotecki et al., 2015; Signorini, Liao, Duncan, Jan, & Stoffel, 1997), and donated by Dr. Kevin Wickman (University of Minnesota). Male *Girk1^flox/flox^* mice were bred with female mice purchased from Jackson Laboratories expressing cre recombinase in parvalbumin-expressing neurons (B6.129P2-Pvalbtm^1(cre)Arbr^/J; Stock No:017320) then back crossed to create *Girk1^flox/flox^*. Female *Girk1^flox/flox^* mice positive for PVcre were then bred with male mice purchased from Jackson Laboratories expressing tdtomato in parvalbumin positive neurons (C57BL/6-Tg(Pvalb-tdtomato)15Gfng/J; Stock No: 027395) and back crossed to generate experimental *Girk1^flox/flox^* mice hemizygous for PVcre and PVtdtomato. Mice were housed in a temperature and humidity-controlled room with a 12h/12h light/dark cycle with food and water available *ad libitum* except throughout attention set-shifting and the progressive ratio test. Male and female experimental mice were PD78 ± 2 days at the start of visual cue testing. Behavioral procedures were conducted in the light phase. Experiments were approved by the Institutional Animal Care and Use Committee at Marquette University.

### Behavioral Testing Timeline

Mice were handled for three days then tested in the elevated plus maze (EPM), followed by attention set-shifting training and testing, progressive ratio, and forced swim test (Figure 1A). A subset of mice were then used for slice electrophysiology, however not all mice used for slice electrophysiology underwent behavioral assessments.

**Figure 1.**
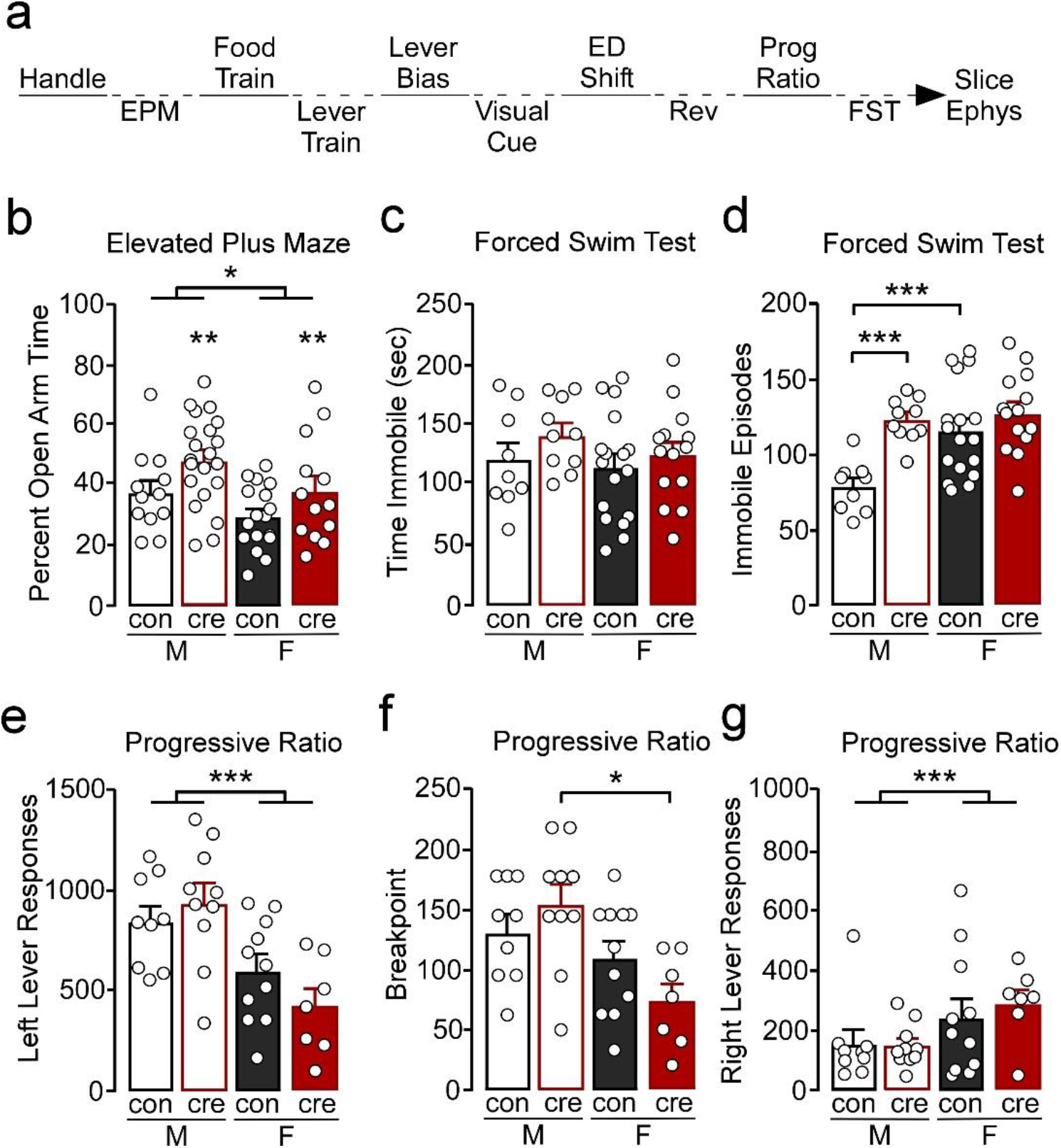
**a)** Experimental timeline. Mice were handled for at least three days then tested in the elevated plus maze (EPM), food and lever trained, followed by assessment for a lever bias. Mice were then tested in a visual cue test, an extradimensional shift (ED shift), and a reversal (REV) test, followed by assessment of motivation in a progressive ratio (Prog Ratio) test. Mice were then tested in the forced swim test (FST). Whole-cell slice electrophysiology was conducted in a subset of mice however not all mice that had slice electrophysiology underwent behavior. **b)** Males spent a greater percent of time in the open arm compared to females (*p<0.05). Cre-positive mice had a significant increase in percent open arm time (**p<0.01). **c)** Male and female control and cre-positive mice had similar amount of immobile time in the forced swim test. **d)** Female control mice had greater number of immobile episodes compared to male control mice (***p<0.001). Male cre-positive mice had greater number of immobile episodes compared to control males (***p<0.001). **e)** During progressive ratio, males had greater number of left lever active presses compared to females (***p<0.001). **f)** Female cre-positive mice had a significantly lower breakpoint compared to male cre-positive mice (*p<0.05). **g)** Female mice had a greater number of right lever inactive presses compared to males, regardless of condition (***p<0.001).

### Elevated Plus Maze

Mice were tested for anxiety-like behaviors using the EPM previously described (Anderson, Gomez, Caccamise, McPhail, & Hearing, 2019) under low light conditions (50 lux at center of maze). AnyMaze (Stoelting Co.) tracking software was used to record and analyze behavior. Percent time in the open arms was calculated as total time in the open arms divided by total time in the maze.

### Attention Set-shifting

Attention set-shifting training and tests were conducted in operant conditioning chambers (Med Associates, Inc.) and were modified and based off methods previously described (Brady & Floresco, 2015). Following EPM, mice were food deprived 85-90% of their free feeding weight. For all attention set-shifting training and testing, correct responses were rewarded with presentation of 50% liquid Ensure^®^ diluted in tap water. During *food training*, a fixed ratio 1 schedule was used, whereby a response on either the left or the right lever resulted in delivery of a reward. The initial food training session was three hours in length and repeated daily until the mouse earned at least 50 rewards. The next day, mice had 30 minutes to receive at least 50 rewards and this training session was also repeated daily until mice did so.

During *lever training*, retractable levers were pseudorandomly presented with no more than two consecutive extensions of each lever. Mice were required to press the lever within 10 seconds of lever extension to receive a reward; absence of a lever press was deemed as an omission. Each lever training session consisted of 90 trials (45 of each the left and right lever), with each trial followed by a 20 second time out. Mice were required to reach criteriion ≤5 omissions total on two consecutive days to move forward. Once lever training criterion was reached, *lever bias* was assessed during which both the right and left lever were presented and reinforced on a fixed ratio 1 for a total of seven trials. If a mouse did not have ≤5 omissions for two consecutive days within 15 days of lever training, training was ceased, and the mouse did not progress through testing (N=12).

The following day, *visual cue testing* was conducted until 150 trials or 10 consecutive correct responses was reached with a minimum of at least 30 trials. If this criteria was not reached on the first day of testing, a second or third day of testing was conducted. During visual cue testing, an illuminated cue light was presented above either the left or right lever in a pseudorandomized order. A response on the lever underneath the illuminated cue light resulted in reward delivery and a 20-second time out, incorrect responses resulted in only the time out. Omissions were counted as stated above and were not counted towards a trial to criteria (i.e. neither correct nor error) when data was analyzed but counted towards the daily 150 trials. The next day, *extradimensional shift testing* was conducted, during which the reinforced lever was always the opposite lever of the previously assessed lever bias. The cue light was presented in a manner/order similar to that during the visual cue test however was not associated with the correct response. Mice were run in the extradimensional shift test until 10 consecutive responses or 150 trials were conducted with a minimum of 30 trials. The day after the extradimensional shift test criteria was reached (i.e. 10 consecutive correct responses), reversal testing was conducted during which the reinforced lever was always that of the previously assessed lever bias, however the cue light was presented identical to that used during the visual cue test. For set-shifting experiments, all tests for a mouse were excluded if >15 omissions in a given test were recorded as this may reflect mechanical issues, motoric, or motivational issues and may influence responding on subsequent tests.

For further behavioral assessment, the types of errors were analyzed based on previously published methods (Brady & Floresco, 2015). Tests were divided into blocks of 16 trials, not including trials were there was omitted responses. For visual cue testing, *initial errors* were errors that were made within each block until there were less than six in a single block. Once there were less than six errors in a single block, errors in all subsequent blocks were characterized as *regressive errors*. For the extradimensional set shift test, tests were divided into bins of 16 trials. Errors that were made based on the previous visual cue test rule used such that the response was made on the lever under the illuminated cue light were tagged and were considered *perseverative* until less than six errors in a single bin were made. Errors in the next bin and subsequent bins were considered regressive errors. Never reinforced errors were those that were incorrect but the response was not on the lever underneath the illuminated cue light. For the reversal test, perseverative and regressive errors were assessed as described above, however errors were considered perseverative until less than 10 errors in a bin were made. Separately, errors were divided into errors that were made towards the cue light distractor and away from the cue light distractor. Response latencies were measured as the amount of time from the extension of the lever until a response was made.

### Progressive Ratio

Following attention set-shifting, mice were then tested for motivation using liquid Ensure^®^ as the reinforcer. Responses on the left lever were reinforced whereas responses on the right lever were inactive and resulted in no consequences, with responses required to obtain each subsequent reward progressively increased. The schedule of reinforcement was (5e^.2*n^)-5 (Richardson & Roberts, 1996) and testing lasted for a total of 90 minutes. Following progressive ratio testing, food was returned *ad libitum*.

### Forced Swim Test

A transparent glass beaker 7” in diameter was filled with 25 ± 2 °C to a depth to prevent the mouse from touching the bottom. Mice were individually placed in the water and habituated for two minutes. Behavioral assessment was recording during the subsequent four minutes during which immobile time and number of episodes were tracked. Following testing, mice were immediately dried and kept in a warm holding cage. Behaviors were recorded using a side mounted camera and assessed using AnyMaze tracking software. Immobility sensitivity was set at 85% and a minimum of 250ms to be counted as an immobility episode.

### Slice Electrophysiology

Mice were anesthetized with isoflurane (Henry Schein), decapitated, and the brain removed and put in ice-cold 95% O_2_ 5% CO_2_ oxygenated sucrose solution (229 mM sucrose, 1.9 mM KCl, 1.2 mM NaH_2_PO_4_, 33 mM NaHCO_3_, 10 mM glucose, 0.4 mM ascorbic acid, 6mM MgCl_2_, and 0.5 mM CaCl_2_) oxygenated using 95% O_2_ 5% CO_2_. Coronal slices (300μm) containing the mPFC were collected using a Leica VT1000S vibratome (Leica VT1000S). Slices were immediately incubated at 31°C for 10 minutes in a solution containing 119mM NaCl, 2.5mM KCl, 1mM NaH_2_PO_4_, 26.2mM NaHCO_3_, 11mM glucose, 0.4mM ascorbic acid, 4mM MgCl_2_, and 1mM CaCl_2_. Slices were then removed, allowed to cool to room temperature and incubated further for a minimum of 35 minutes.

Whole-cell recordings were performed as previously described (Anderson et al., 2019; Hearing et al., 2013). Oxygenated ACSF was gravity perfused at a temperature of 29°C-33°C using at a flow rate of ~2-2.5 ml/minute. Sutter Integrated Patch Amplifier (IPA) with Igor Pro (Wave Metrics, Inc.) was used for the data acquisition software. Recordings were filtered at 2kHz and sampled at 5kHz. Layer 5/6 PVI were identified based on the presence of td-tomato fluorescence. For all recordings, adequate whole-cell access (Ra<40 MΩ) was maintained. Borosilicate glass pipettes were filled with 140mM K-Gluconate, 5.0mM HEPES, 1.1mM EGTA, 2.0mM MgCl_2_, 2.0mM Na_2_-ATP, 0.3mM Na-GTP, and 5.0mM phosphocreatine (pH 7.3, 290mOsm). For rheobase and action potential frequency, a 20pA current-step injection was used (0-300pA). For ML297 recordings, a baseline with <20% fluctuation current was obtained, followed by bath application of 10μM ML297 (David Weaver, Vanderbilt) in 0.04% DMSO (Sigma-Aldrich). Evoked currents were reversed using bath application of 0.30mM barium chloride (Fisher Scientific). Sample sizes are denoted as *n* for the number of recordings/cells and *N* for the number of mice.

### Statistical Analysis

Data are presented as mean ± SEM. SigmaPlot 11.0 was used to perform statistical analyses. A 2 (male, female) x 2 (control, cre) analysis of variance were used for all comparisons except assessment of progressive ratio break point which violates parametric assumptions and therefore a Kruskal-Wallis test was used. Student-Newman-Keuls method for multiple post-hoc comparisons was used when applicable. Statistical outliers (±2SD) from behavioral tests were excluded from analyses (2 data points from EPM; 5 total mice from all attention set-shift tests; 1 data point from progressive ratio).

## RESULTS

### Effects of GIRK1 knockout in PVI on EPM, forced swim test, and motivation

Constitutive knockout of GIRK signaling, including channels expressing the GIRK1 subunit, has been shown to alter anxiety- and depression-like behavior, learning and memory, as well as motivation for appetitive rewards (Llamosas, Bruzos-Cidon, Rodriguez, Ugedo, & Torrecilla, 2015; Pravetoni & Wickman, 2008; Victoria et al., 2016; Wydeven et al., 2014). However, recent work has shown that cell-type specific ablation of GIRK signaling has unique effects on behavior (Victoria et al., 2016), making straightforward interpretation of these phenotypes difficult. As GIRK channels are known to be present in PVI and alteration of PV-dependent GABA neuron activity is known to regulate affect and cognitive control (D. Kim et al., 2016; Murray et al., 2015; Page, Shepard, Heslin, & Coutellier, 2019; Rossi et al., 2012; Sohal et al., 2009; Sparta et al., 2014), we aimed to determine whether GIRK1-dependent signaling in PVI is necessary for normal affect, motivation, and cognitive control.

To examine the behavioral relevance of PVI GIRK1 signaling, we first assessed affect-related behavior using the EPM and FST. Individual differences in locomotor activity were controlled for by assessing the percent of time spent in the open arm divided by the total time assessed. A main effect of sex was identified, with males having increased percent time in the open arm compared to females (F_(1,59)_= 5.91, p=0.018). There was also a main effect of condition, with cre-positive mice having increased percent open arm time compared to control mice (F_(1,59)_= 7.05, p=0.010) however no sex by condition interaction was observed (F_(1,59)_= 0.07, p=0.789; Figure 1b).

During the FST, there were no differences in the total time spent immobile (sex: F_(1,46)_= 0.87, p=0.356; condition: F_(1,46)_= 1.91, p=0.174; interaction: F_(1,46)_= 0.20, p=0.657; Figure 1c). There was however, a significant sex by condition interaction when comparing number of immobile episodes during testing (F_(1,46)_= 4.78, p=0.034), with male cre-positive mice having significantly more immobile episodes than control counterparts (p<0.001) whereas there were no differences between conditions in female mice (p=0.200). Female controls also had increased immobile episodes compared to the male controls (p<0.001) whereas male and female cre-positive mice did not differ (p=0.619; Figure 1d).

To examine if loss of GIRK1 in PVI impacts motivation and to control for any potential differences observed in our set-shifting experiments (see below), we assessed responding for an appetitive liquid reward (Ensure^®^) using a progressive ratio model. Responses on the left reinforced lever, the breaking point at which the mouse would no longer respond, and the right nonreinforced lever were recorded. Males had an overall greater number of left lever presses compared to females, regardless of condition, whereas controls and cre-positive mice had similar number of responses (sex: F_(1,33)_= 18.50, p<0.001; condition: F_(1,33)_= 0.22, p=0.640; interaction: F_(1,33)_= 2.43, p=0.128; Figure 1e). A Kruskal-Wallis nonparametric test detected a significant difference on break point during progressive ratio (H_(3)_= 10.76, p=0.013) with break points in male cre-positive greater than female cre-positive mice, with no other significant post-hoc comparisons (Figure 1f). Notably, similar to the left lever, a main effect of sex was observed for responding on the right non-reinforced lever (F_(1,33)_= 5.44, p=0.026). In this case females displayed a higher number of presses, however no effect of condition or interaction was observed (condition: F_(1,33)_= 0.15, p=0.702; interaction: F_(1,33)_= 0.26, p=0.614; Figure 1g).

### Effects of PVI GIRK1 knockout on cognitive flexibility

Our recent work has shown that disruption of GIRK1 signaling in mPFC prelimbic pyramidal neurons impairs cognitive performance in an attentional set-shifting model of cognitive flexibility (Anderson et al., 2020; biorxiv). Given prior research highlighting a role for PVI activity in cognitive control, we next assessed whether GIRK1 signaling in PVI also influences cognitive control. Prior to attention set-shift testing, mice had to have two consecutive days with less than or equal to five omissions during lever training during which there was no effect of sex (F_(1,34)_= 0.94, p=0.340), condition (F_(1,34)_= 1.64, p=0.210), nor a interaction (F_(1,34)_= 1.59, p=0.216; data not shown). Once mice passed lever training criterion and were assessed for a side bias, testing in the visual cue was conducted. There was an effect of sex on the total trials to reach criteria during the visual cue test (F_(1,34)_= 8.32, p=0.007) but no effect of condition (F_(1,34)_= 0.42, p=0.522) or a sex by condition interaction (F_(1,34)_= 0.41, p=0.527; Figure 2a). Similarly, there was an effect of sex on the total errors made to reach criteria during the visual cue test (F_(1,34)_= 8.32, p=0.007) but no effect of condition (F_(1,34)_= 0.31, p=0.583) nor a sex by condition interaction (F_(1,34)_= 0.41, p=0.710; Figure 2b).

**Figure 2.**
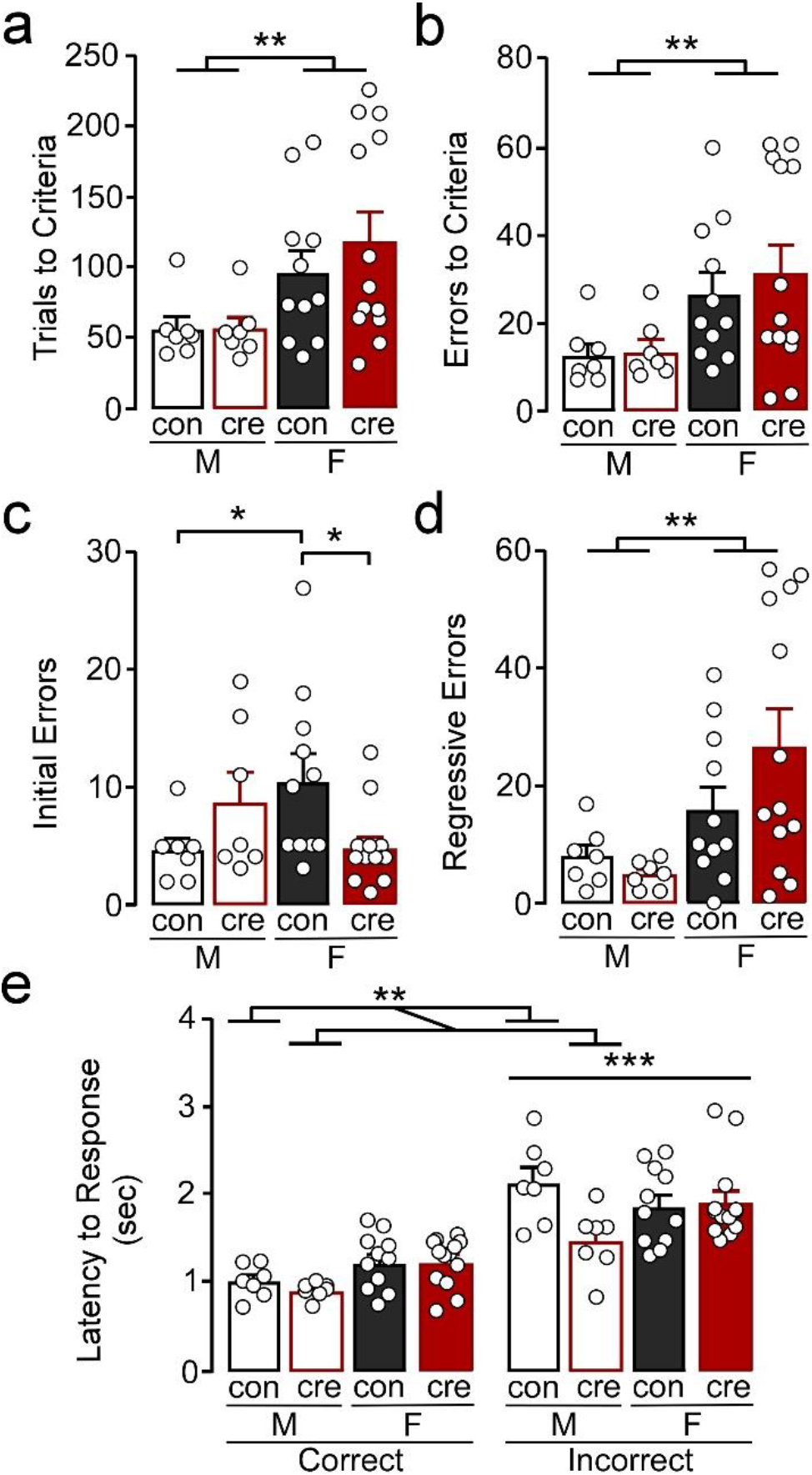
Visual cue test. **a)** Females took more trials to reach criteria compared to males (**p<0.01). **b)** Females made more errors to reach criteria compared to males (**p<0.01). **c)** Female control mice made more initial errors compared to male control mice while female cre-positive made less errors compared to female control mice (*p<0.05). **d)** Females made more regressive errors compared to males (**p<0.01). **e)** The latency to make an incorrect response was higher than the latency to make a correct response (***p<0.001). Cre-positive male mice, regardless of response type, took less time to respond compared to control mice (**p<0.01).

In a more refined investigation into errors, we divided error type based on errors made prior to when mice received less than six errors in one 16 trial bin (i.e. initial errors) and those after (regressive errors), the latter of which reflects an inability to maintain the rule strategy. For initial errors there was a significant sex by condition interaction (F_(1,34)_= 7.72, p=0.009; Figure 2c) with post-hoc comparisons indicating no significant difference between male control and cre-positive mice (p=0.154) whereas female cre-positive mice had fewer initial errors compared to control mice (p=0.012). Notably, control females had significantly more initial errors compared to control males (p=0.027) whereas male and female cre-positive mice did not differ (p=0.116). Examination of regressive errors showed no main effect of condition (F_(1,34)_= 0.61, p=0.441) or a condition by sex interaction (F_(1,34)_=1.98, p=0.169), however females had significantly more regressive errors compared to males (F_(1,34)_= 9.09, p=0.005; Figure 2d). Together, further analyses of the error type reveals that regressive errors drive the increased errors to criteria in females while assessment of the initial errors indicate female control mice have greater difficulty at the beginning of the test compared to male controls and female cre-positive mice.

The speed of processing and general motor function of each mouse can be measured by taking the average latency to respond after the lever was presented (Brady & Floresco, 2015). A 2 (sex) x 2 (condition) x 2 (response type) ANOVA was used to compare latency to response during the visual cue test. There was a main effect of response type with latency for mice to respond being significantly greater for incorrect compared to correct responses (F_(1,68)_=76.88, p<0.001; Figure 2e). There was also a significant sex by condition interaction (F_(1,68)_= 6.01, p=0.017) with post-hoc comparisons indicating that male cre-positive mice took less time to respond compared to male controls (p=0.006) while females regardless of condition had similar response latencies (p=0.758). There were no other significant interactions (sex by response type: F_(1,68)_= 1.06, p=0.307; condition by response type: F_(1,68)_= 2.09, p=0.153; sex by condition by response type: F_(1,68)_= 3.06, p=0.085).

During the extradimensional shift test, there were no differences in trials to reach criteria comparing sex (F_(1,34)_= 0.84, p=0.365), condition (F_(1,34)_= 0.21, p=0.653), or sex by condition (F_(1,34)_= 0.00, p=0.978; Figure 3a). There were also no differences in errors to reach criteria comparing sex (F_(1,34)_= 2.22, p=0.145), condition (F_(1,34)_= 0.79, p=0.381), or sex by condition (F_(1,34)_= 0.37, p=0.551; Figure 3b). However, females had overall more perseverative errors than males (F_(1,34)_=4.34, p=0.045) while there were no differences of condition on number of perseverative errors (F_(1,34)_= 0.71, p=0.404) or a sex by condition interaction (F_(1,34)_=0.03, p=0.882; Figure 3c). There was no main effect of sex (F_(1,34)_= 0.72, p=0.402), main effect of condition (F_(1,34)_= 0.01, p=0.910), nor a sex by condition interaction (F_(1,34)_= 0.01, p=0.094) of regressive errors during the extradimensional shift test (Figure 3d). There were also no differences on number of errors made that had previously never been reinforced (sex: F_(1,34)_= 2.27, p=0.141; condition: F_(1,34)_= 0.03, p=0.857; interaction: F_(1,34)_= 1.28, p=0.265; Figure 3e). There was a significant effect of sex on latency to respond, regardless of condition and response type, with females taking longer to respond compared to males (F_(1,68)_= 7.39, p=0.008; Figure 3f) but no other differences (condition: F_(1,68)_= 0.21, p=0.645; response type: F_(1,68)_= 0.08, p=0.772; sex by condition: F_(1,68)_= 0.00, p=0.974; sex by response type: F_(1,68)_= 0.00, p=0.973; condition by response type: F_(1,68)_= 0.64, p=0.427; sex by condition by response type: F_(1,68)_= 0.26, p=0.614).

**Figure 3.**
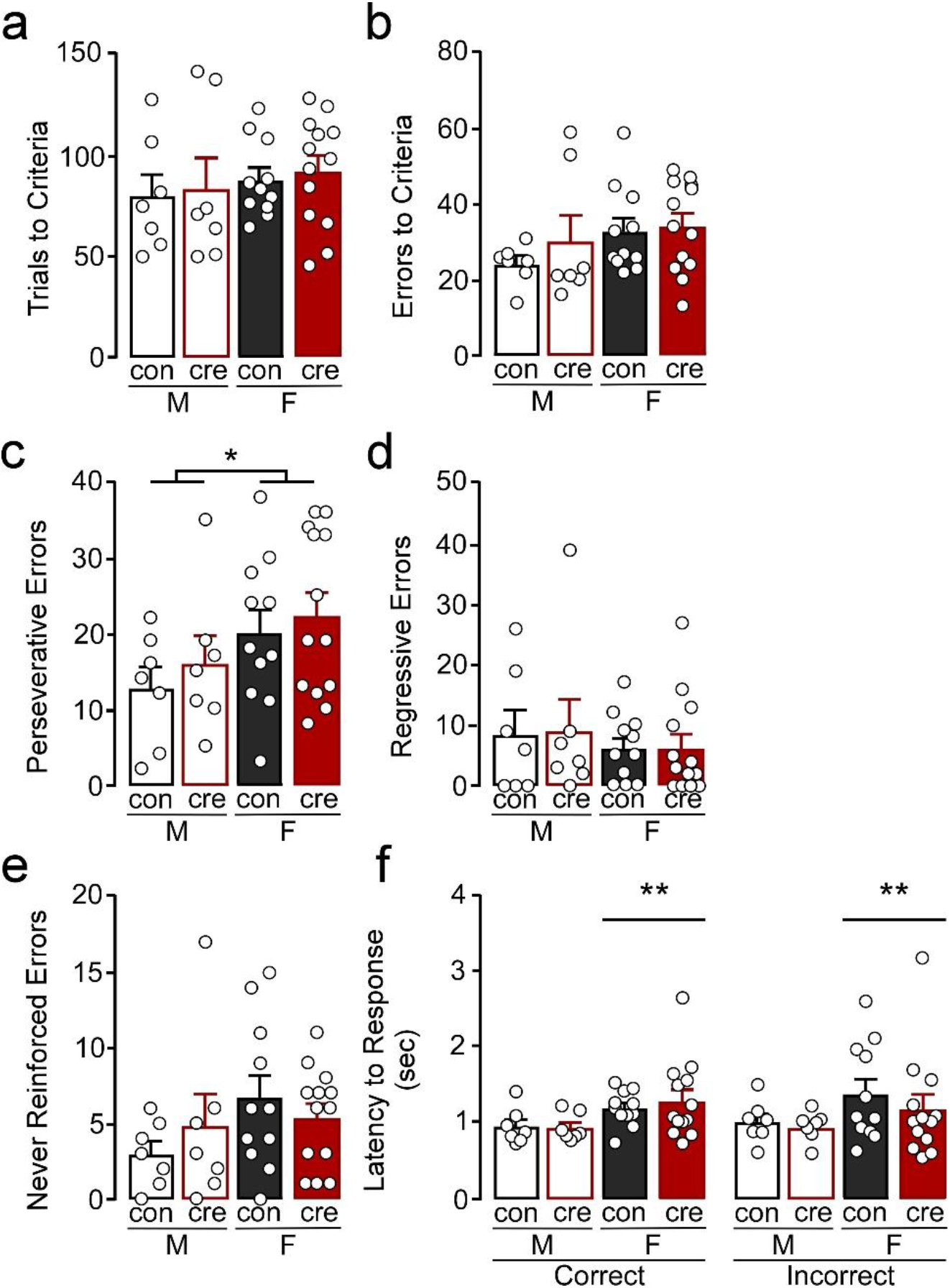
Extradimensional shift test. **a)** Trials to criteria and **b)** errors to criteria were similar for all groups. **c)** Females made more perseverative errors compared to males (*p<0.05) while the number of **d)** regressive and **e)** never reinforced errors were similar for all groups. **f)** Females took longer to respond compared to males (**p<0.01) regardless of condition and response type.

During the reversal test, females took more trials to reach criteria compared to males (F_(1,33)_= 8.03, p=0.008) but there were no differences comparing conditions (F_(1,33)_=1.59, p=0.216) or a sex by condition interaction (F_(1,33)_= 0.04, p=0.849; Figure 4a). Similarly, females had more errors during the reversal test compared to males (F_(1,33)_=4.54, p=0.041), however there was no difference comparing the conditions (F_(1,33)_=0.24, p=0.630) or a sex by condition interaction (F_(1,33)_=0.07, p=0.793; Figure 4b). There were no differences in perseverative errors (sex: F_(1,33)_= 0.16, p=0.695; condition: F_(1,33)_= 0.02, p=0.882; sex by condition: F_(1,33)_= 0.00, p=0.969; Figure 4c) or regressive errors (sex: F_(1,33)_= 3.94, p=0.056; condition: F_(1,33)_= 0.12, p=0.727; sex by condition: F_(1,33)_= 0.14, p=0.709; Figure 4d). Errors were next analyzed as being either toward or away from the cue distractor. There was a significant effect of sex on errors towards the distractor (F_(1,33)_=5.30, p=0.028) with females making more errors towards the distractor (Figure 4e) but there were no differences between the two conditions (F_(1,33)_= 1.11, p=0.299) or a sex by condition interaction (F_(1,33)_=0.00, p=0.988). There were also no differences in number of errors made away from the distractor (sex: F_(1,33)_= 1.43, p=0.241; condition: F_(1,33)_= 0.24, p=0.629; interaction: F_(1,33)_= 0.31, p=0.582; Figure 4f). Finally, there were no differences in latency to make a response during the reversal test (sex: F_(1,66)_= 2.17, p=0.146; condition: F_(1,66)_= 1.81, p=0.184; response type: F_(1,66)_= 0.44, p=0.510; sex by condition: F_(1,66)_= 1.00, p=0.322; sex by response type: F_(1,66)_= 0.45, p=0.506; condition by response type: F_(1,66)_= 0.00, p=0.944; sex by condition by response type: F_(1,66)_= 0.79, p=0.377; Figure 4g). The extradimensional shift and reversal tests suggest that cre-positive mice have similar cognitive flexibility to control mice and also reveal that females tend to persist on the initial visual cue test rule (i.e. increased perseverative errors during the extradimensional shift test and increased errors towards distractor during the reversal test) despite the changing contingencies.

**Figure 4.**
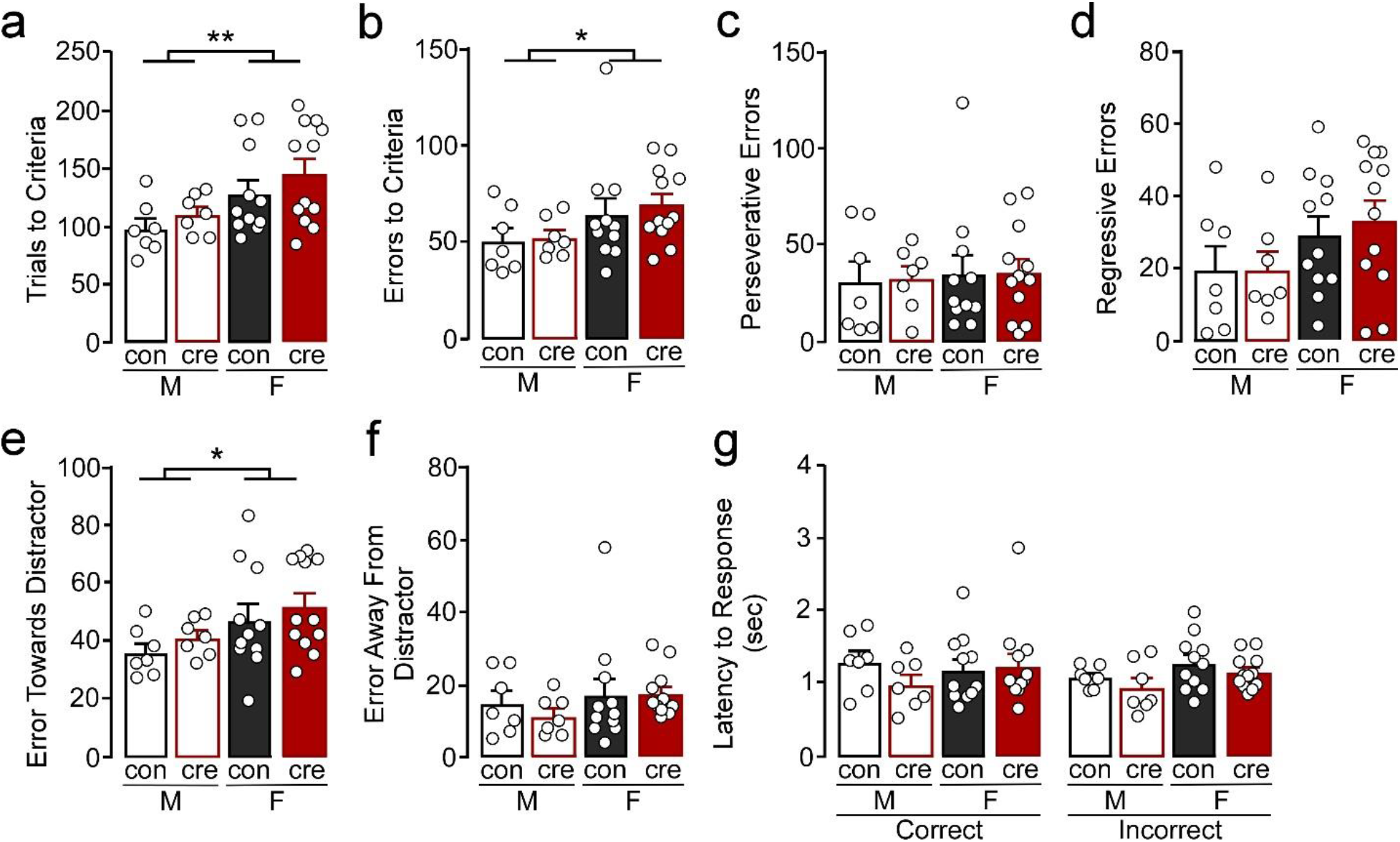
Reversal test. **a)** Females took more trials to criteria (**p<0.01) and **b)** errors to criteria (*p<0.01) compared to males. **c)** All groups had similar number of perseverative and **d)** regressive errors. **e)** Females made more errors towards the cue distractor compared to males (*p<0.05). **f)** The number of errors away from the distractor was similar for all groups. **g)** All groups had similar latency to respond, regardless of response type.

### Characterization of GIRK1 knockout in prelimbic cortex PVI

Past work has identified an important role for GIRK1 signaling in hippocampal PVI (Booker et al., 2013). Given known contributions of the prelimbic region of the mPFC in regulating the aforementioned behaviors, we confirmed the presence of GIRK1 in prelimbic PVI and characterized knockout in *Girk1^flox/flox^* mice. To facilitate the electrophysiological evaluation of PVI neurons in PVCre: *Girk1^flox/flox^* mice, we crossed this line with transgenic mice expressing tdtomato under the control of the PV promoter. To determine if the knockout was reducing GIRK1 in PVI, voltage-clamped whole-cell slice recordings were performed to assess GIRK1-specific changes in somatodendritic currents using the GIRK1 selective agonist, ML297 (Wydeven et al., 2014)(Figure 5a). Bath application of ML297 produced an outward current (I_ML297_) that correlated with a decrease in input resistance (not shown) and was blocked by subsequent application of barium chloride (0.3 mM). Comparison across sex and condition (con vs. cre) showed that ML297-mediated currents were not different in PVI from males versus females (F_(1,21)_= 0.26, p=0.616) nor was there a sex by condition interaction (F_(1,21)_= 0.00, p=0.972). PVI recorded from cre-positive mice showed a significant reduction in ML297-induced current compared to PVI from control mice (F_(1,21)_= 10.23, p=0.004; Figure 5b) indicative of a reduction in GIRK1 signaling (male control n=6/N=4; male cre n=8/N=4; female control n=6/N=5; female cre n=5/N=3).

**Figure 5.**
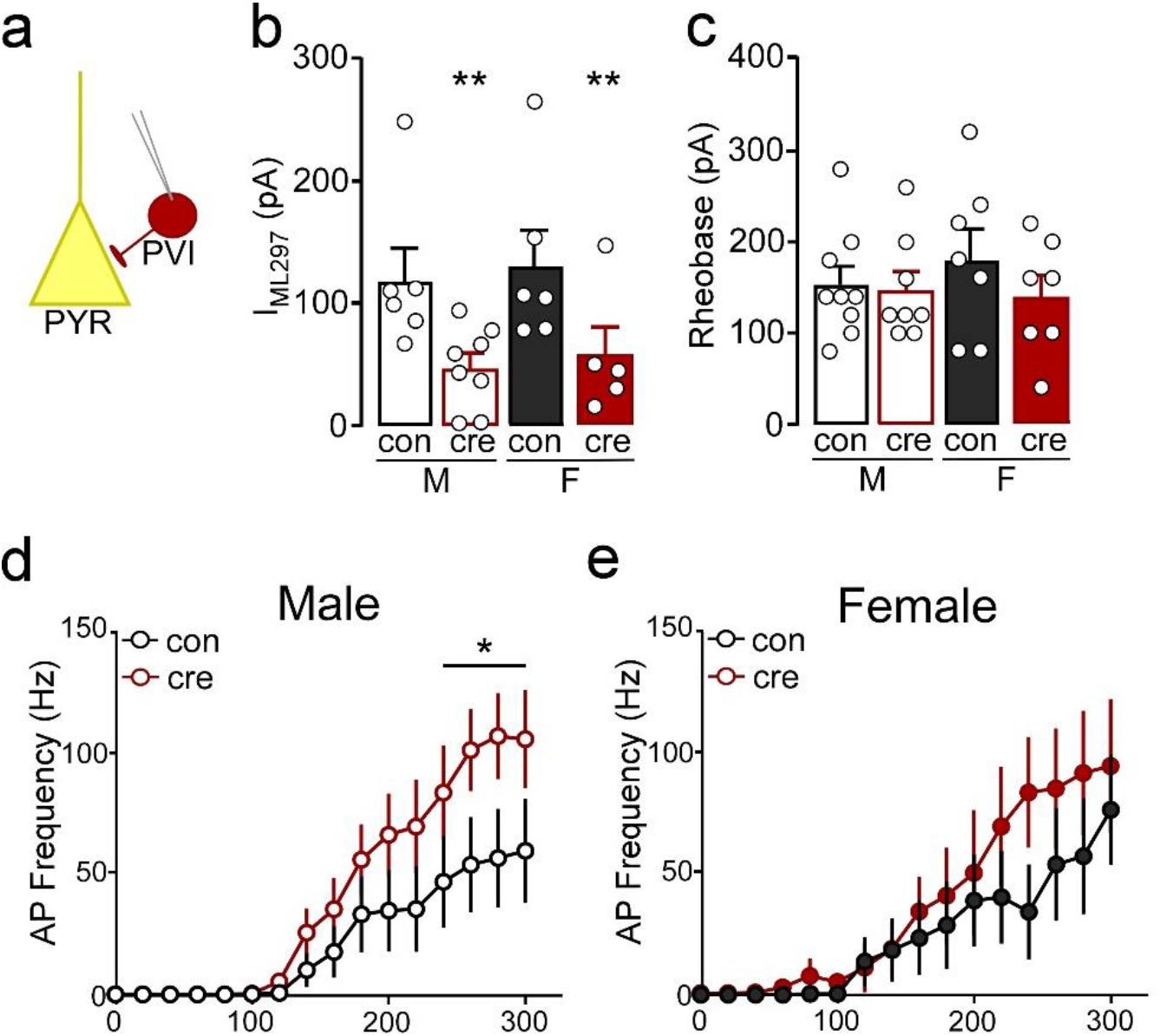
**a)** Whole-cell slice electrophysiology was used to record from parvalbumin interneurons (PVI) labelled with td-tomato in the prelimbic cortex. **b)** The current induced by bath application of ML297 was lower in PVI from cre-positive mice compared to those from control mice (**p<0.01). **c)** The current required to evoke an action potential was similar for all groups. **d)** The number of action potentials fired at 240-300pA was higher in PVI from cre-positive male mice compared to control male mice (*p<0.05). **e)** PVI had similar action potential frequency in cre-positive and control female mice.

To determine how loss of GIRK1 signaling impacts PVI cell membrane excitability, we examined the threshold to fire an action potential (rheobase) and spike firing frequency in response to increasing current injections (male control n=9/N=6; male cre n=8/N=4; female control n=7/N=4; female cre n=7/N=4). Rheobase did not differ based on sex or condition (sex: (F_(1,27)_= 0.21, p=0.651; condition: F_(1,27)_= 1.02, p=0.322; interaction: (F_(1,27)_= 0.59, p=0.449; Figure 5c). During the current-step injection in PVI from males, there was a significant current by condition interaction (F_(15,225)_= 2.01, p=0.016) with cre-positive cells firing significantly more action potentials at higher currents (240-300pA) compared to controls (p<0.05; Figure 5d). Conversely, there was no main effect of condition (F_(1,12)_= 0.69, p=0.423) or a condition by current interaction (F_(15,180)_= 0.75, p=0.735; Figure 5e) in female PVI. Together, these data indicate that loss of GIRK1 does not impact activation threshold but does increase neuronal firing of male PVI.

## DISCUSSION

The current study evaluated the impact of selectively ablating GIRK channels expressing the GIRK1 subunit in PVI on affect and cognitive flexibility. We found that loss of PVI GIRK1 signaling in males and females increased percent time spent in the open arm during the EPM suggestive of reduced anxiety-related behavior. Notably, effects on anxiety-like behavior in EPM or open field were not previously observed with conditional knockout of GIRK2 channels in GABA neurons using GAD-Cre transgenic mice (Victoria et al., 2016), suggesting that more selective targeting of GABA neuron subpopulations may yield discrete changes. Loss of GIRK1 selectively in males also increased immobile episodes during the FST, suggesting an increase in active coping. Although not directly comparable, these findings are in alignment with recent work showing that increased activation of mPFC PVI results in anxiety-like behaviors (Page et al., 2019). Moreover, the lack of effect on motivation during the progressive ratio is in agreement with previous research showing that optogenetic stimulation of mPFC PVI does not alter appetitive reward consumption (Sparta et al., 2014). Past *in vivo* work has shown that increased spike firing of PVI cells reduces output of pyramidal neurons in the mPFC (Sparta et al., 2014), thus it is possible that any observed changes in behavior, albeit discrete, reflect reductions in prelimbic cortex pyramidal neuron activity.

Given the span of literature highlighting a critical role for PVI in regulation of mPFC-dependent cortical processing (Ferguson & Gao, 2018) and previously identified roles for GABA_B_ and GIRK signaling in regulating affect and cognition (Cooper et al., 2012; Lazary et al., 2011; Lecca et al., 2016; Pravetoni & Wickman, 2008; Victoria et al., 2016; Wydeven et al., 2014; Yamada et al., 2012), we predicted that effects of GIRK1 ablation would be evident in measures of cognitive flexibility. These predictions were based on studies using approaches that reduced PVI-dependent signaling, however the impact of increased PVI output on cortical processing and cognition is far less clear. For example, cre-inducible channelrhodopsin activation of mPFC PVI at high frequencies has been shown to promote delayed alternation impairments (Rossi et al., 2012) and accelerate extinction of cue-reward behavior (Sparta et al., 2014). Conversely, chemogenetic and optogenetic activation of mPFC PVI does not alter performance in a novel object recognition test of working memory or fear conditioning (Page et al., 2019; Yizhar et al., 2011). In the present study, loss of PVI GIRK1 signaling did not result in behavioral deficits during overall performance in tests of visual cue-based discriminative learning or flexibility during the extradimensional shift and reversal test – tasks which have been shown to be dependent on the mPFC (Floresco, Block, & Tse, 2008; Ragozzino, Detrick, & Kesner, 1999) and orbitofrontal cortex (Ghods-Sharifi, Haluk, & Floresco, 2008), respectively. However, during the visual cue test, cre-positive males had a reduced latency to respond compared to their control counterparts, which may be indicative of increased speed of processing. These findings align with a reduction in attentional processing following PVI silencing (H. Kim et al., 2016). Although the alterations in speed of processing did not correspond to differences in the trials or errors to criteria, others have shown that PVI inactivation produces deficits in cognitive flexibility in the water maze (Murray et al., 2015). The outcomes of these studies are difficult to compare to the current study, as they determined the role of acute and intermittent increases in PVI activity, rather than chronically altering PVI activity specifically through reductions in GIRK signaling. Regardless, the present study in combination with past work suggest that while loss of PVI signaling promotes critical deficits in cognitive control, the effects of increased PVI output are far more complex. In agreement, a recent report (Caballero, Flores-Barrera, Thomases, & Tseng, 2020) has suggested that there may be a threshold of PVI expression through which cortical dysfunction becomes evident. Thus, it is possible that loss of GIRK signaling alone does not meet that threshold, as other inhibitory signaling may be decreased to compensate for changes in PVI excitability and output.

### GIRK1 knockout effects on PVI excitability

Past studies have shown that GABA_B_-type receptors regulate perisomatic inhibition of hippocampal PVI and that this inhibition is likely driven by activation of GIRK channels (Booker et al., 2013). While GIRK1-3 subunits were shown to be present in PVI, GIRK1 was the most prominently expressed. Our past work has shown that GIRK1-expressing GIRK channels are the primary mediators of GIRK signaling in prelimbic cortex pyramidal neurons, however to our knowledge this is the first study to assess the role of GIRK signaling in prelimbic cortex PVI. Whole-cell recordings with the selective GIRK1 agonist ML297 showed that GIRK1-containing channels are indeed present in prelimbic PVI and that the presence of cre-recombinase reduced GIRK1-mediated currents by ~60%, with residual current likely driven by vehicle (DMSO). While GIRK1 knockout produced a leftward shift in current-spike relationship (increased firing frequency) in PVI from males and to a lesser non-significant extent in females, it unexpectedly did not alter rheobase.

The lack of effect on membrane excitability (as well as lack of a robust behavioral phenotype) may reflect a variety of factors. First, the knockout of PVI GIRK-signaling was already in effect during critical development periods which may have led to compensatory changes in non-GIRK effectors more readily modulating membrane excitability. Although a viral vector to specifically target PVI GIRK-signaling is not readily available, it would be beneficial for future research to target PVI GIRK channels specifically during adulthood to reduce the potential for compensation of other signaling during development. Further, neuronal GIRK channels can be homo- and heterotetrameric complexes formed primarily by assembly of GIRK1, GIRK2, and GIRK3 subunits (Karschin et al., 1996; Hearing et al., 2013), it is possible that loss of GIRK1 leads to upregulation of homotetrameric GIRK2-expressing channels. However, such a phenomenon has not been observed with constitutive and conditional knockout of GIRK1 or GIRK2 in other cell-types in areas such as the hippocampus, prefrontal cortex, and ventral tegmental area (Hearing et al., 2013; Kotecki et al., 2015; Victoria et al., 2016). Third, mPFC PVIs include basket and chandelier type cells that differ in structure, physiology, and GPCR agonist response (Booker et al., 2013; Kawaguchi & Kubota, 1997). Thus, decreasing GIRK-signaling in both cell-types may have diminished any opportunity for behavioral changes that may have arisen should GIRK-signaling have been reduced in only one cell-type. Finally, it is possible that while GIRK channels do in fact play a role in regulating membrane excitability in PVI, these GIRK channels exhibit differences in coupling efficiency or regulators of G protein signaling (Labouebe et al., 2007).

### Sex differences in cognitive flexibility and motivation

Although not the main focus of the study, we unexpectedly identified differences in male and female performance across the set-shifting task that were largely independent of GIRK ablation. During the initial visual cue test, females required more trials and errors to reach criteria compared to males, regardless of genotype. In controls, the increased number of errors in females reflects an equal balance between initial and regressive errors. However, loss of PVI GIRK appears to shift this towards an increase in regressive error types and a reduction in initial errors, suggesting an impaired ability to maintain a newly learned rule.

During the extradimensional shift, while females do not require more trials to reach criteria than males, they perseverate more on the rule associated with the prior day visual cue test (they continue to follow the cue light) and take longer to respond compared to males. Females also require more trials to reach criteria in a test of reversal learning. Further investigation of the error types show a greater number of errors on the lever underneath the cue (i.e. more errors towards the distractor) indicating that they are still perseverating on the visual cue rule, despite learning a new rule during the extradimensional shift. To our knowledge, this is the first study to investigate sex differences in an operant based attentional set-shifting model of cognitive flexibility. It should be noted that although gonadal hormones influence cognition (Luine & Frankfurt, 2020; Taxier, Gross, & Frick, 2020), the estrous cycle does not appear to influence attention set-shifting performance (Workman, Crozier, Lieblich, & Galea, 2013), and these sex differences may not be replicated in C57BL/6J mice. The day following the reversal test, motivation for an ensure reward was measured using a progressive ratio test during which only the left lever was reinforced. Although male mice had significantly greater number of left lever responses, females exhibited greater responding on the right lever. As progressive ratio testing only occurred on a single day, the increased right lever responses in females may suggest a lack of task understanding with a shift to right lever responding at higher break points.

Findings from the current study suggest that PVI GIRK1 signaling does not mediate membrane excitability to the extent it does in principle pyramidal neurons and may differentially impact firing frequency in males and females. Further, while reductions in PVI GIRK1 signaling influenced anxiety-like behaviors, effects on cognitive performance were more nuanced. While outside the scope of this study, the demonstration that GIRK channels are present in prelimbic PVI requires further investigation into their function, as past work has shown that differences in subunit expression and coupling dictate responsivity to various Gi-coupled GPCR agonists, and thus may inform future drug therapies targeting GIRKs or Gi-coupled GPCRs. Further, the current findings have provided unexpected insight into how biological sex impacts cognitive processing associated with an operant-based model of cognitive flexibility which may have important implications for treating pathological deficits in cognitive control.

## Acknowledgements

These studies were supported by funding from the Brain and Behavior Research Foundation (#26299), Marquette University Regular Research Grant, and the Charles E Kubly Mental Health Research Foundation at Marquette University.

